## SUPPLEMENTARY INFORMATION for "LinkedSV for detection of mosaic structural variants from linked-read exome and genome sequencing data"

Fang et al.

### Supplementary Figures

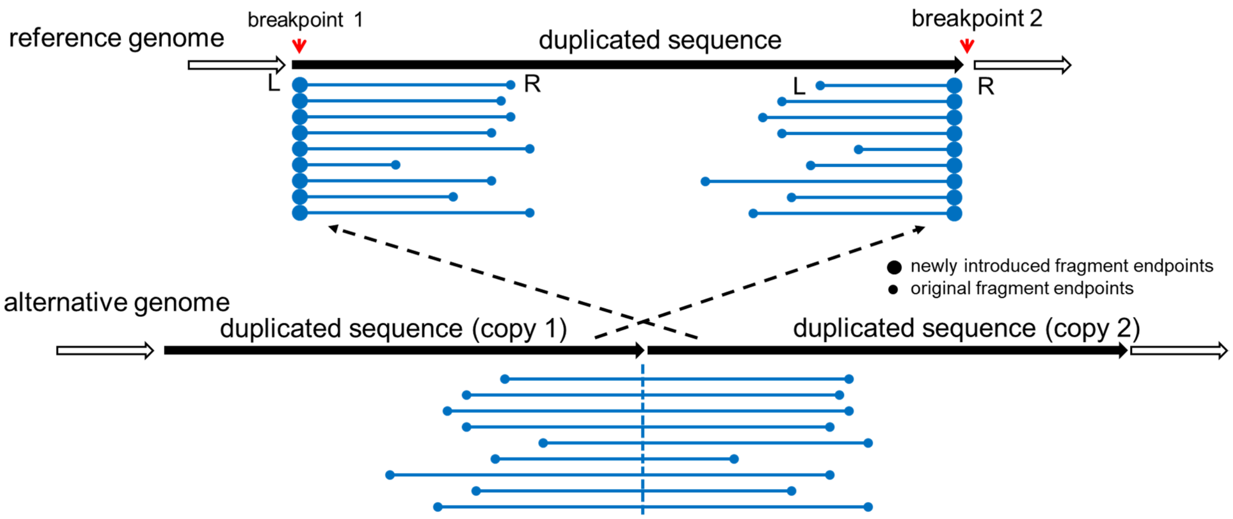


#### Supplementary Figure 1

The pattern of enriched fragment endpoints for tandem duplications. L-endpoints and R-endpoints are enriched near breakpoint 1 and breakpoint 2, respectively. Breakpoints are marked by red arrows.

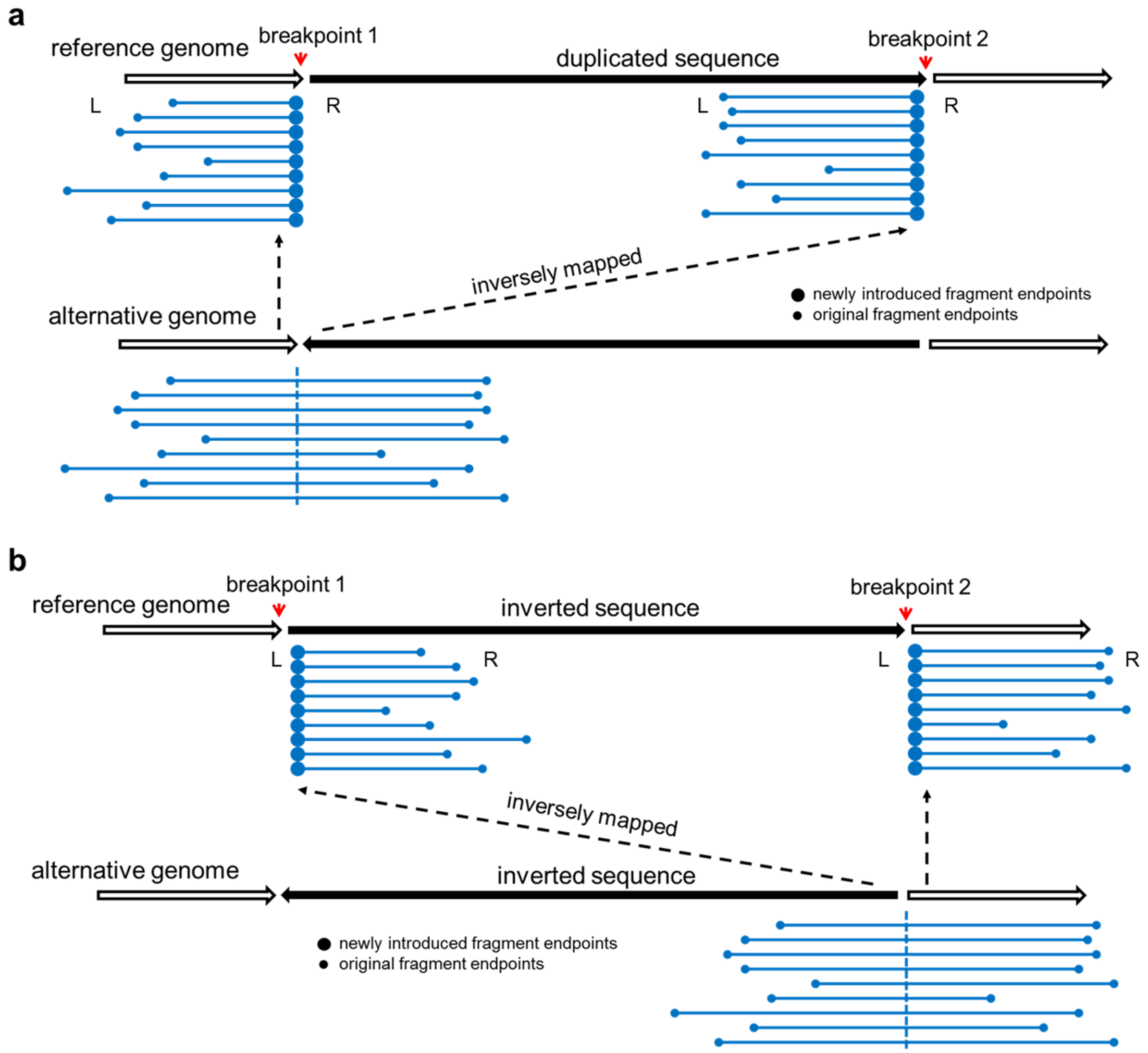


**Supplementary Figure 2**

The pattern of enriched fragment endpoints for inversions. The endpoints enriched near the two breakpoints are of the same type. **a**) Enriched fragment endpoints are both R-endpoints. **b**) Enriched fragment endpoints are both L-endpoints.

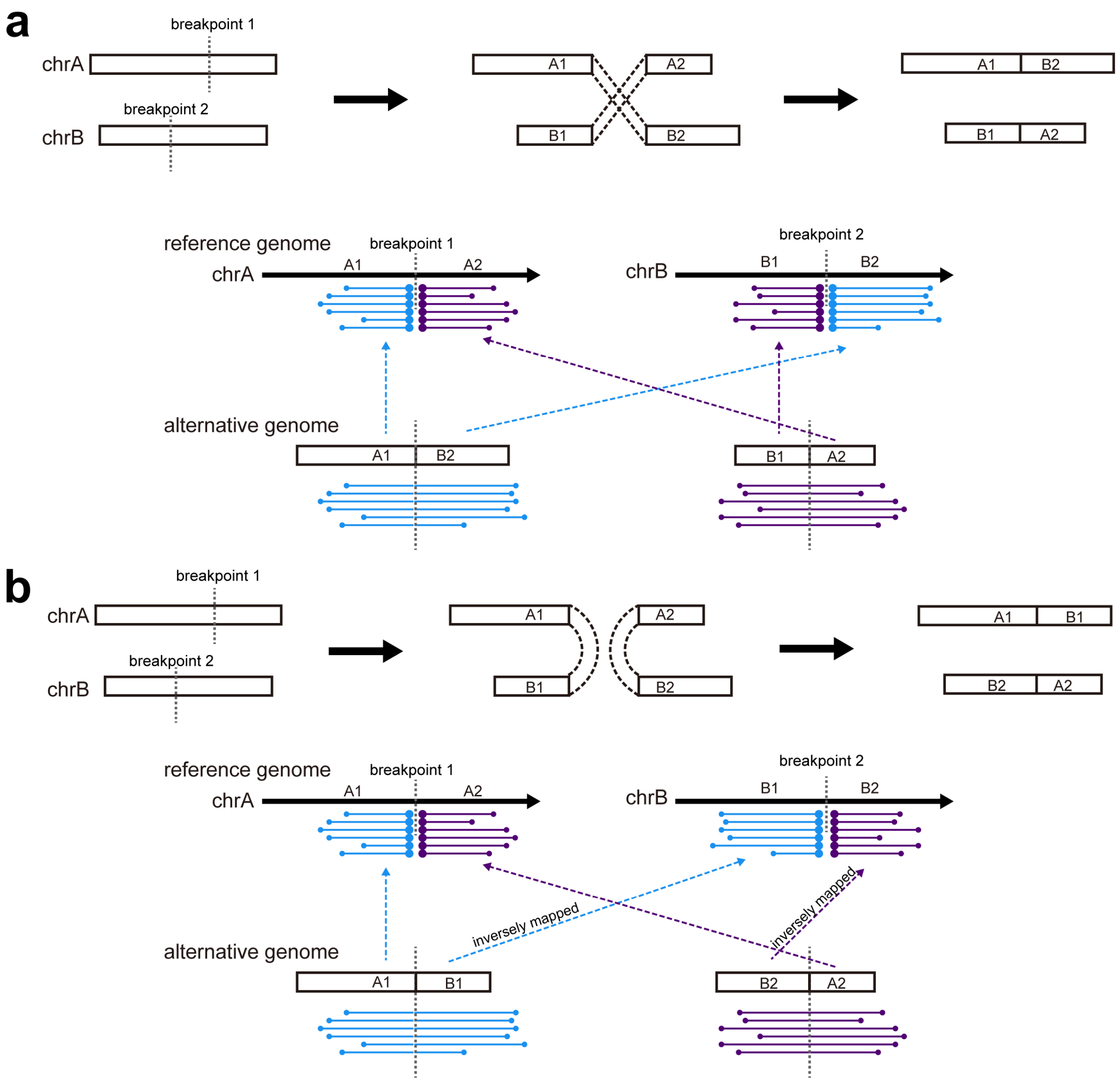


**Supplementary Figure 3**

The pattern of enriched fragment endpoints for interchromosomal translocations. The enriched endpoints can be either L-endpoints or R-endpoints, depending on how the two chromosomes are joined. **a**) two chromosomes are joined in the same direction. **b**) two chromosomes are joined in the reverse direction.

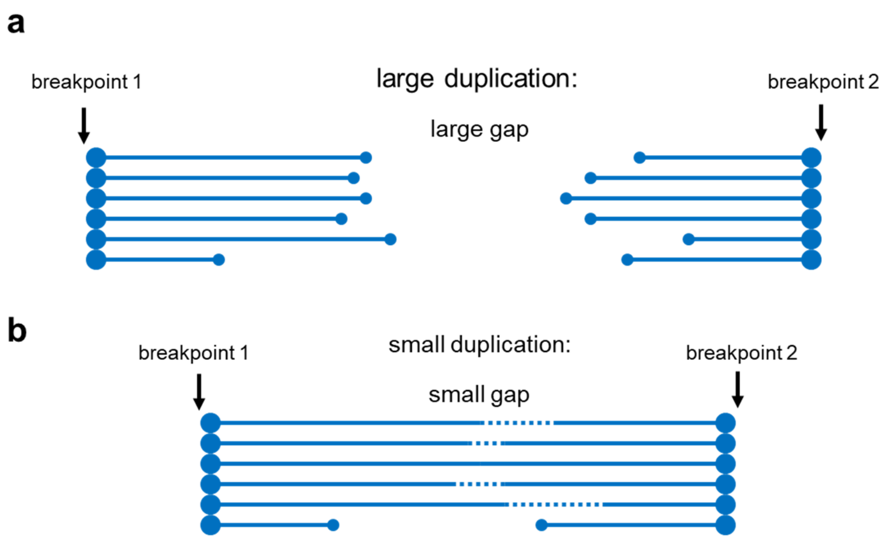


**Supplementary Figure 4**

**a**) For large duplications, the reads of the alternative allele are separated by a large gap so that we can observe two sets of fragments with the same set of barcodes, which indicate an SV. **b**) If the duplication is not large enough, the reads will be probably clustered into one fragment.

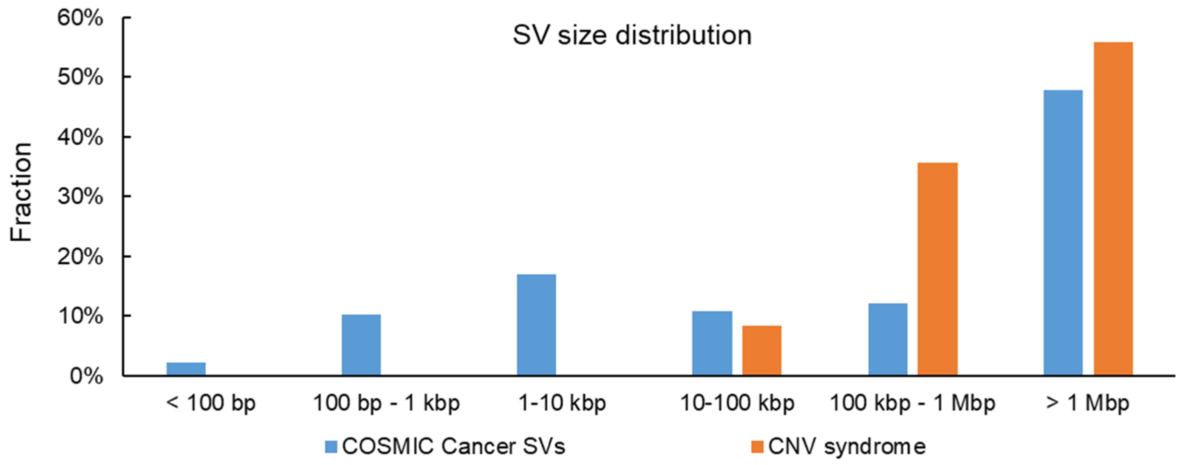


**Supplementary Figure 5**

Size distribution of SVs from two resources: cancer somatic SVs in COSMIC database and expert-curated known CNVs that cause CNV syndromes. Inter-chromosomal events are regarded as > 1Mbp here because in terms of SV detection using linked-reads, the inter-chromosomal events share the same properties with super large intra-chromosomal events. Source data is provided as a Source Data file.

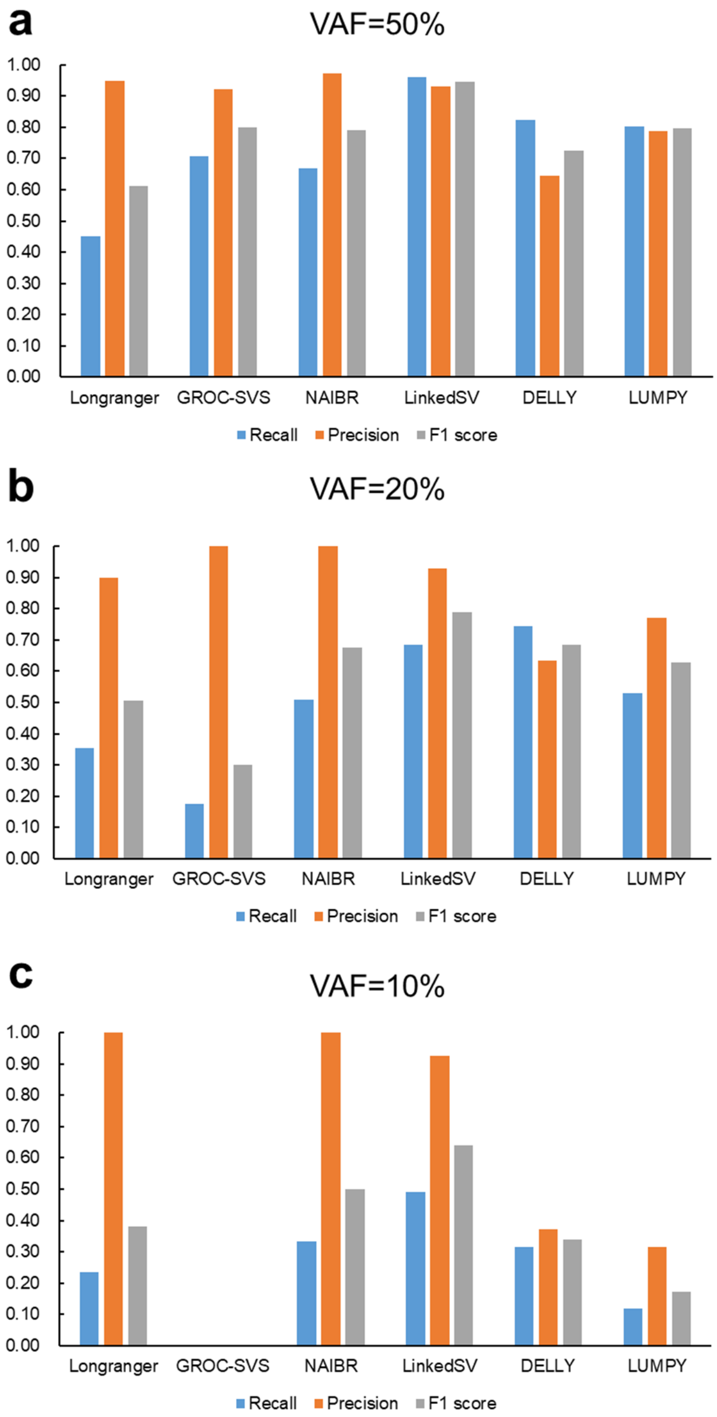


**Supplementary Figure 6**

Benchmarking of six SV callers on simulated deletions and duplications that cause CNV syndromes. **a**) Variant allele frequency (VAF) = 50%. This is a simulation of germline variants. **b**)

VAF = 20%. **c**) VAF = 10%. **b**) and **c**) are the simulations of somatic or mosaic variants. Source data is provided as a Source Data file.

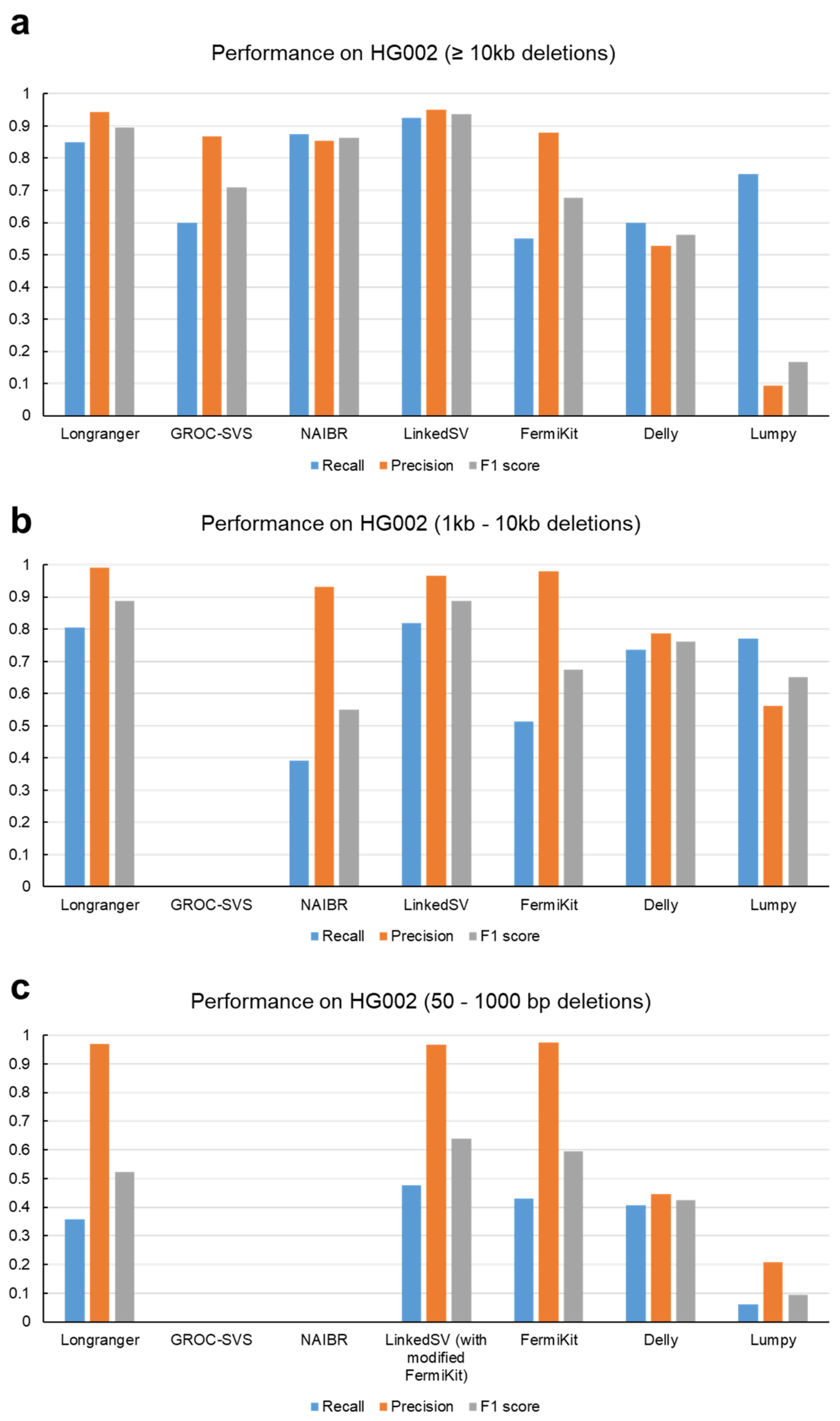


**Supplementary Figure 7**

Benchmarking of deletion detection on the HG002 genome. **a**) Performance of detection of deletions that are > 10 kb. **b**) Performance of detection of deletions that are within 1-10 kb. **c**) Performance of detection of deletions that are within 50-1000 bp. Source data is provided as a Source Data file.

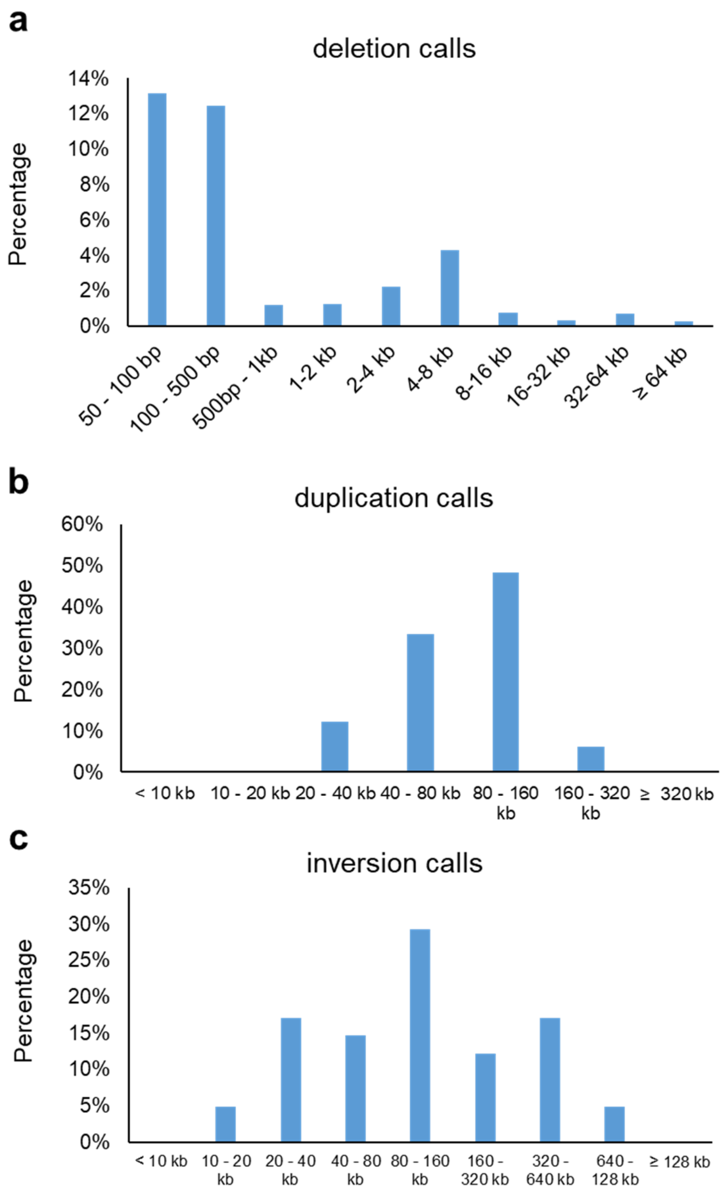


**Supplementary Figure 8**

Size distribution of SV events detected from the HG002 genome using LinkedSV. **a**) Deletions. The small peak in 4-8 kb indicate the events of LINE elements, which are about 7 kb long. **b**)

Duplications. **c**) Inversions. LinkedSV is able to detect small deletions up to 50 bp by using multiple information including paired-end read signals and local assembly (using modified FermiKit). LinkedSV currently only uses barcode information to detect duplications and inversions, therefore, only large duplications and inversion can be detected. Source data is provided as a Source Data file.

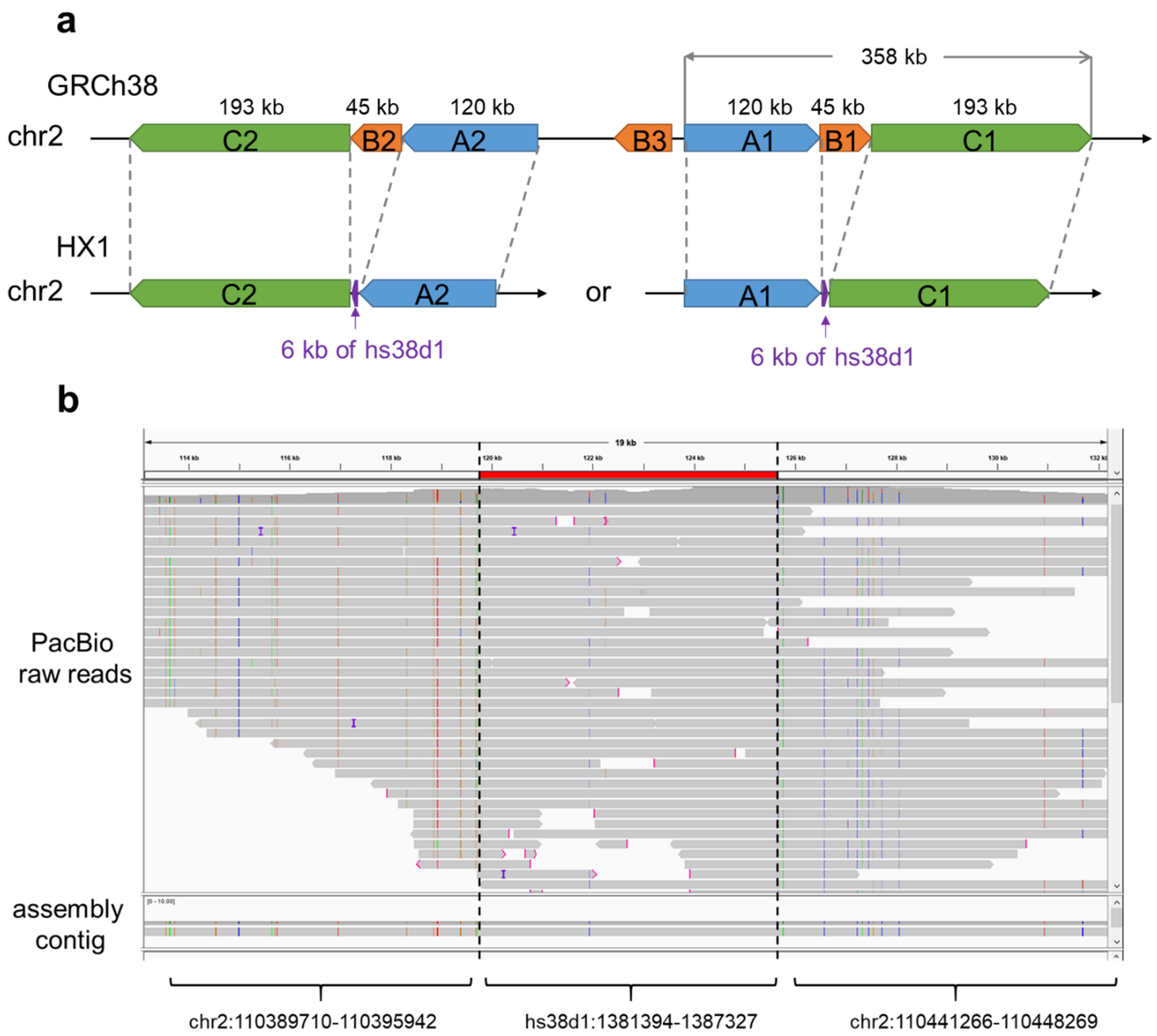


**Supplementary Figure 9**

**a**) Proposed variant allele in HX1 chr2. The 45 kb deletion region resides in a 358 kb segmental duplication region, which has two copies in chr2. The two copies are in the opposite direction and are highly identical (fraction of matched bases = 99.88%, according to UCSC genome browser). The 45 kb deletion region has a third copy between the two 358 kb segmental duplications. In the HX1 genome, the 45 kb region is deleted and a 6 kb region from hs38d1 decoy sequence is inserted. Since the two 358 kb segmental duplications are large and highly identical, with the current read length and sequencing error rate, we are unable to tell which copy contains this event. **b**) Alignment of PacBio raw reads as well as the assembly of the aligned reads to the proposed variant allele sequence (A1 + 6 kb insertion + C1). Reads with mapping quality > 20 were shown.

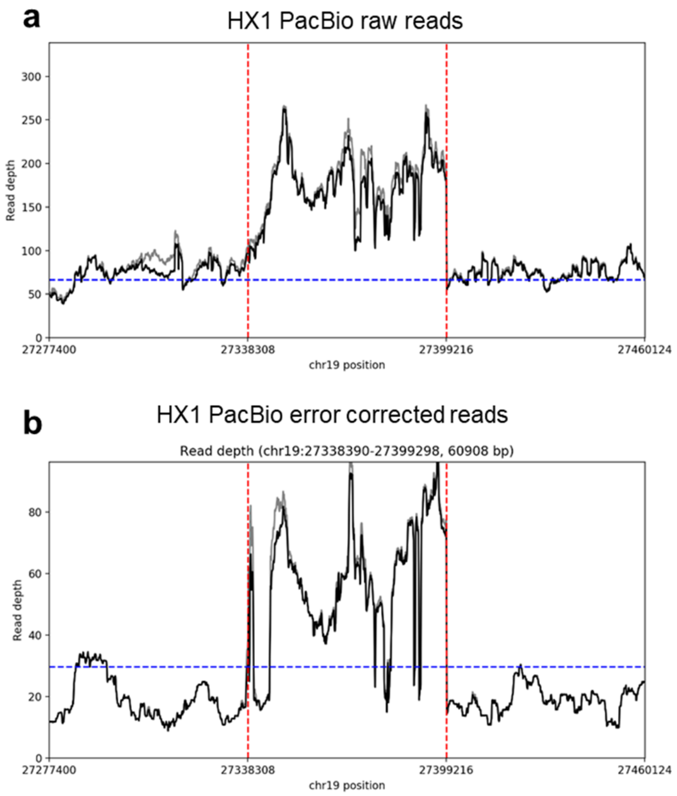


**Supplementary Figure 10**

Sequencing coverage of HX1 raw reads (**a**) and error corrected reads (**b**) near the chr19 duplication region detected by LinkedSV. Sequencing coverage of the linked reads is shown in Figure 7d. The predicted breakpoints by LinkedSV were indicated by vertical red lines. The dotted blue line showed the average depth across the whole genome.

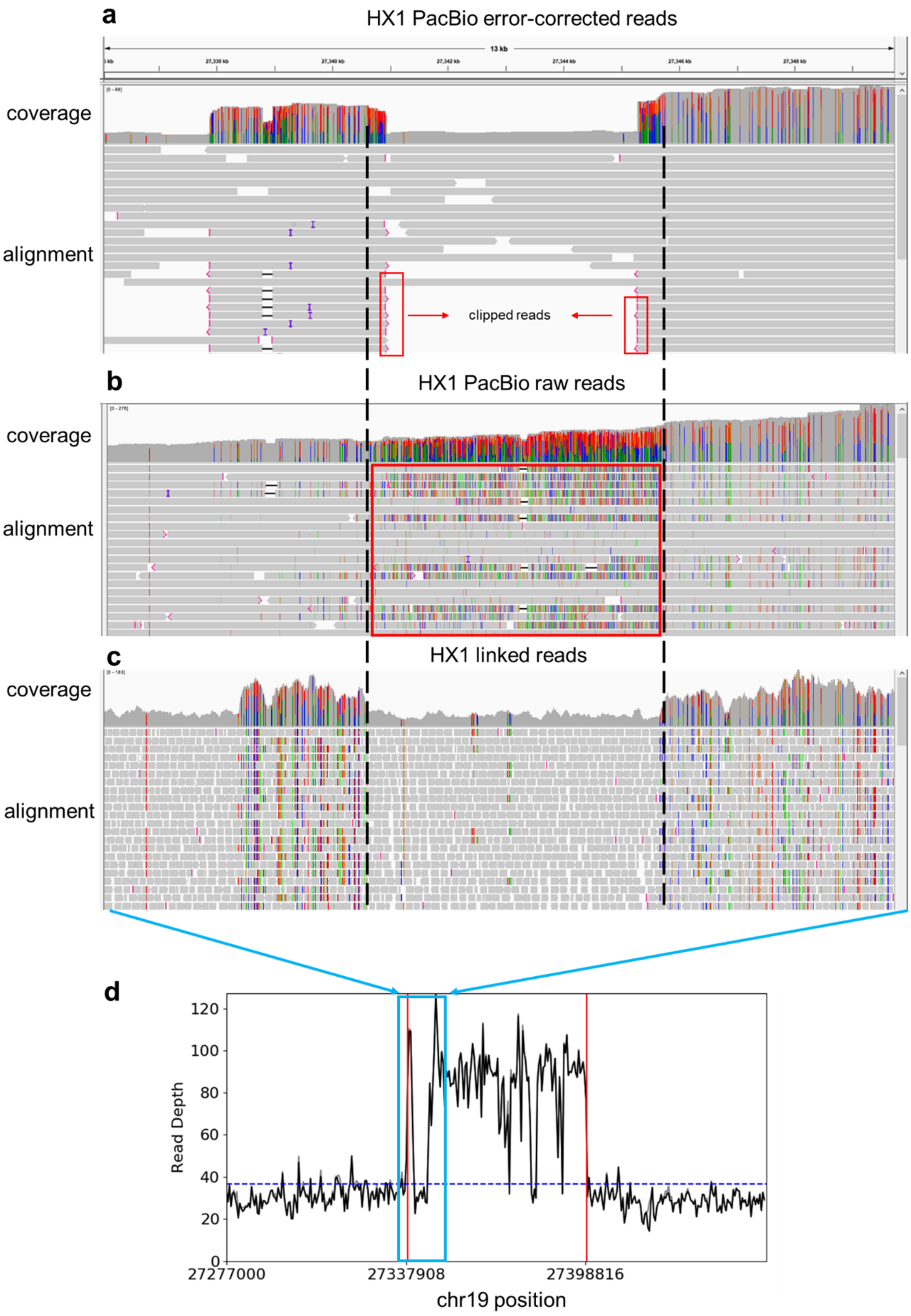
a
HX1 PacBio error-corrected reads
13 kb
coverage
alignment
clipped reads
b
HX1 PacBio raw reads
coverage
alignment
c
HX1 linked reads
coverage
alignment
d
Read Depth
0
20
40
60
80
100
120
27277000
27337908
27398816
chr19 position

**Supplementary Figure 11**

Inspection of the start location of the chr19 duplication in HX1 rewiews two duplication events. **a**) Sequencing coverage and alignments of error-corrected PacBio reads, as shown by IGV. A small duplication event was found next to the main event. From the coverage track, clear boundaries of the two events can be seen. In the alignment track, the clipped reads were marked by pink lines (5'-clipping) or pink arrows (3'-clipping). The alignments were genrated by minimap2 with parameters for PacBio read (-x map-pb) **b**) Sequencing coverage and alignments of raw PacBio reads, as shown by IGV. In the alignment track, mismatch bases were shown in colors (A, green; T, red; C, blue; G, orange). There are enriched alignment mismatch in the red box, indicating that this portion of reads should be clipped, rather than aligned. This may explain that the boundaris of the two events were not clear in the the coverage track. The alignments were genrated by minimap2 with parameters for PacBio read (-x map-pb). **c**) Sequencing coverage and alignments of 10X Genomics linked reads, as shown by IGV. In the coverage track, the boundaries of the two events can be seen and they are consitent with error-corrected PacBio reads. **d**) Zoom out view of the whole duplication region. Y-axis shows the sequencing coverage of 10X Genomics linked reads. The dotted blue line showed the average depth across the whole genome. The predicted breakpoints by LinkedSV were indicated by vertical red lines. This pannel is the same as Figure 7d.

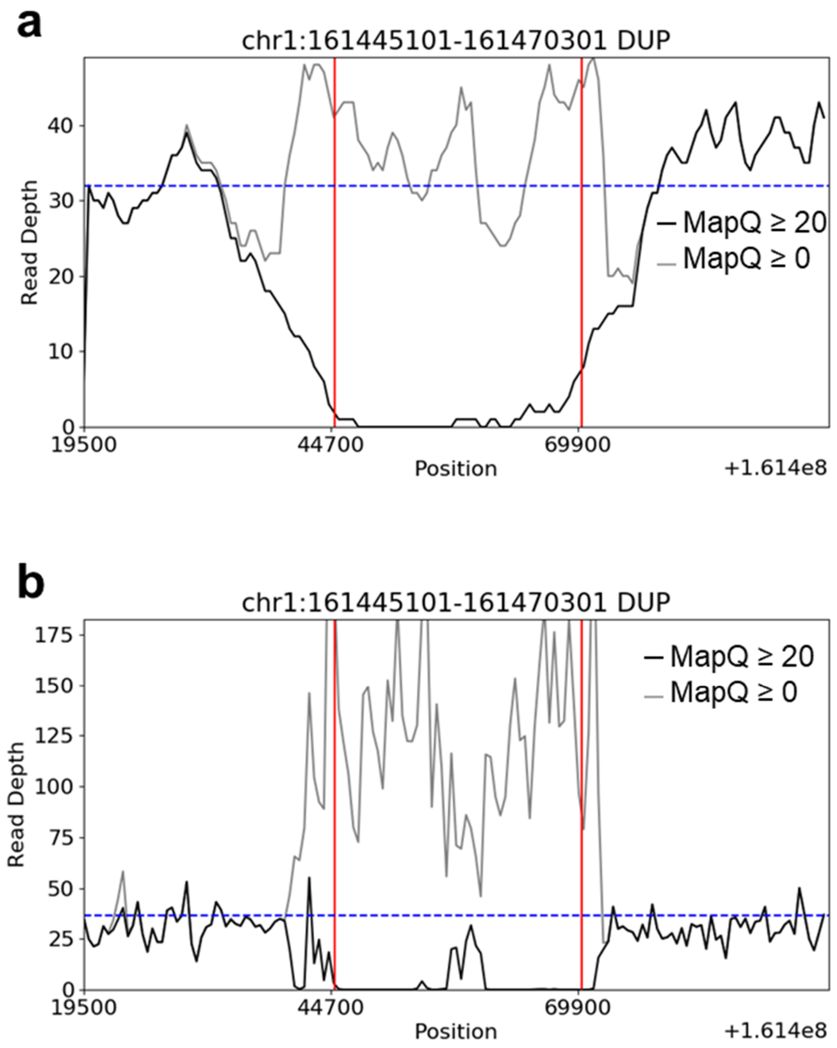


**Supplementary Figure 12**

Read depth distribution of PacBio long reads (**a**) and linked reads (**b**) near a duplication reported by SMRT-SV. The black lines showed the read depth of reads with mapping quality ≥ 20 while the grey lines showed the read depth of reads with mapping quality ≥ 0 (i.e. all reads). The read depth was calculated using SAMtools. The dotted blue line showed the average depth across the whole genome. The large space between the black line and the grey line indicates poor mapping qualities in the predicted duplication region.

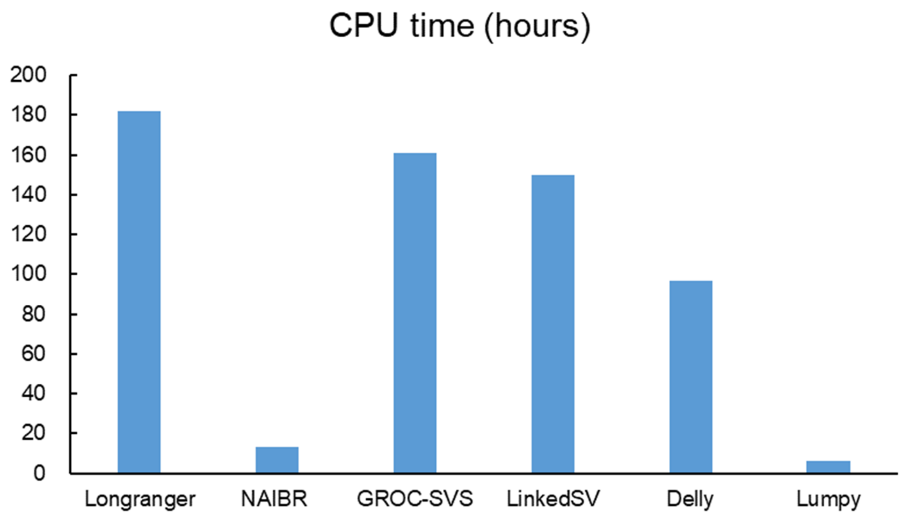


**Supplementary Figure 13**

Computation time (in hours) of different SV callers on the 37X coverage HX1 WGS data set. LinkedSV uses longer time than NAIBR, Delly and Lumpy, because it uses two types of barcode evidence and also performed local assembly to detect small deletions. Source data is provided as a Source Data file.

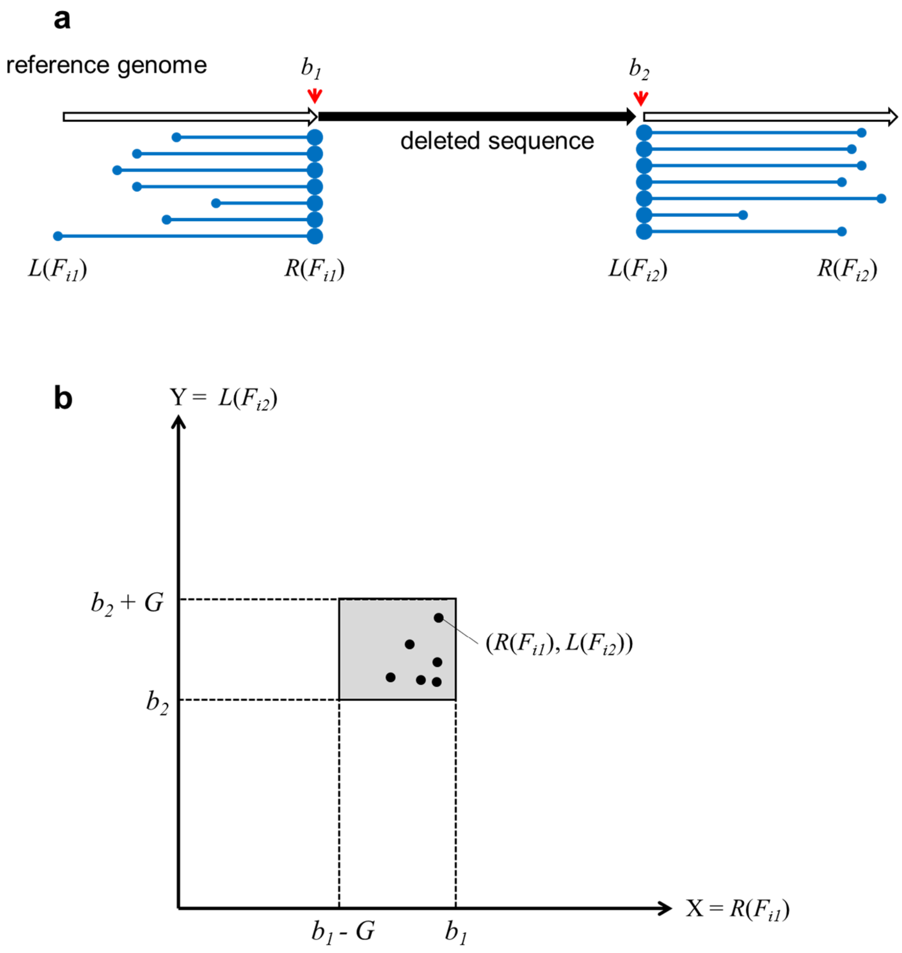


**Supplementary Figure 14**

Detection of type 1 evidence. We use deletion as an example but the method is also suitable for the other SV types. **a**) R-endpoints ($R(F_{i1})$) and L-endpoints ($L(F_{i2})$) are enriched near the deletion breakpoints $b_1$ and $b_2$, respectively. **b**) Two-dimensional plot of $R(F_{i1})$ and $L(F_{i2})$. $R(F_{i1})$ and $L(F_{i2})$ are restricted in the grey square according to equation (2). The background noise of the two-dimensional plot is cleaner than the one-dimensional plot since the fragments that do not share barcodes are excluded.

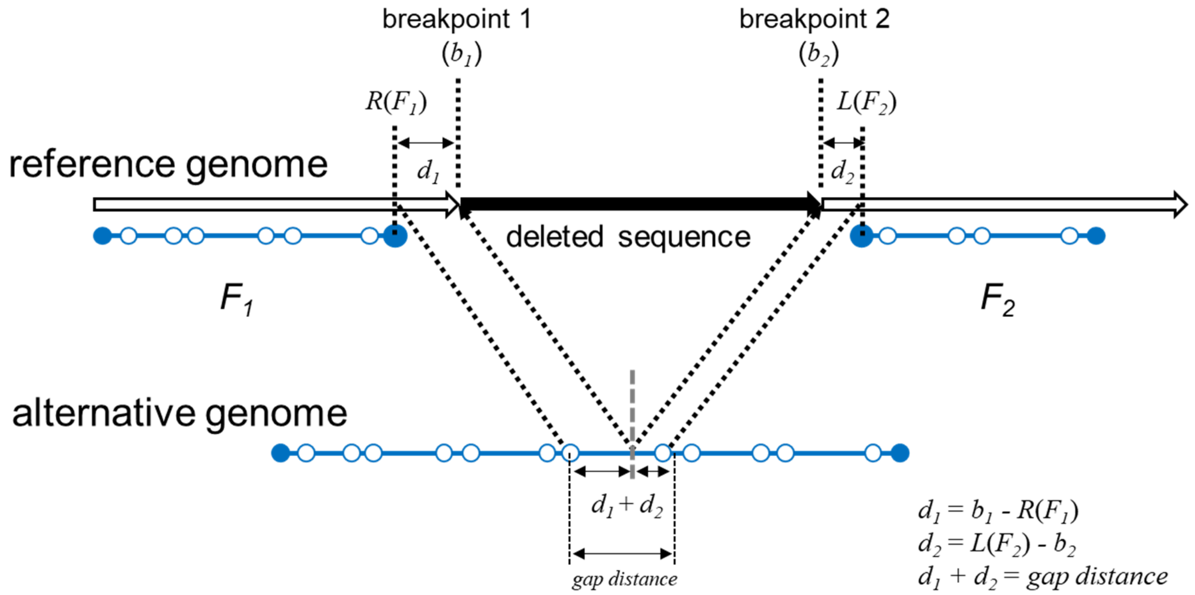


**Supplementary Figure 15**

Explanation of the enrichment of fragment endpoints near the breakpoints, using a deletion as an example. In this event, reads from a HMW DNA molecule that spans the breakpoints of a deletion were mapped to two genomic locations, resulting in two observed fragments (denoted by $F_1$ and $F_2$). $b_1$ and $b_2$ denote the positions of breakpoint 1 and 2. $R(F_1)$ denotes the right endpoint of $F_1$ and $L(F_2)$ denotes the left endpoint of $F_2$. $d_1$ and $d_2$ denote the distances between the fragment endpoints ($R(F_1)$, $L(F_2)$) and the corresponding breakpoints ($b_1$, $b_2$), respectively. In the original fragment of the alternative genome, $d_1 + d_2$ equals to the distance between two adjacent reads (i.e. gap distance). Therefore, gap distance is the upper limit of $d_1$ and $d_2$. Solid blue dots are endpoints of the fragments. Hollow circles are the short reads in the fragments.

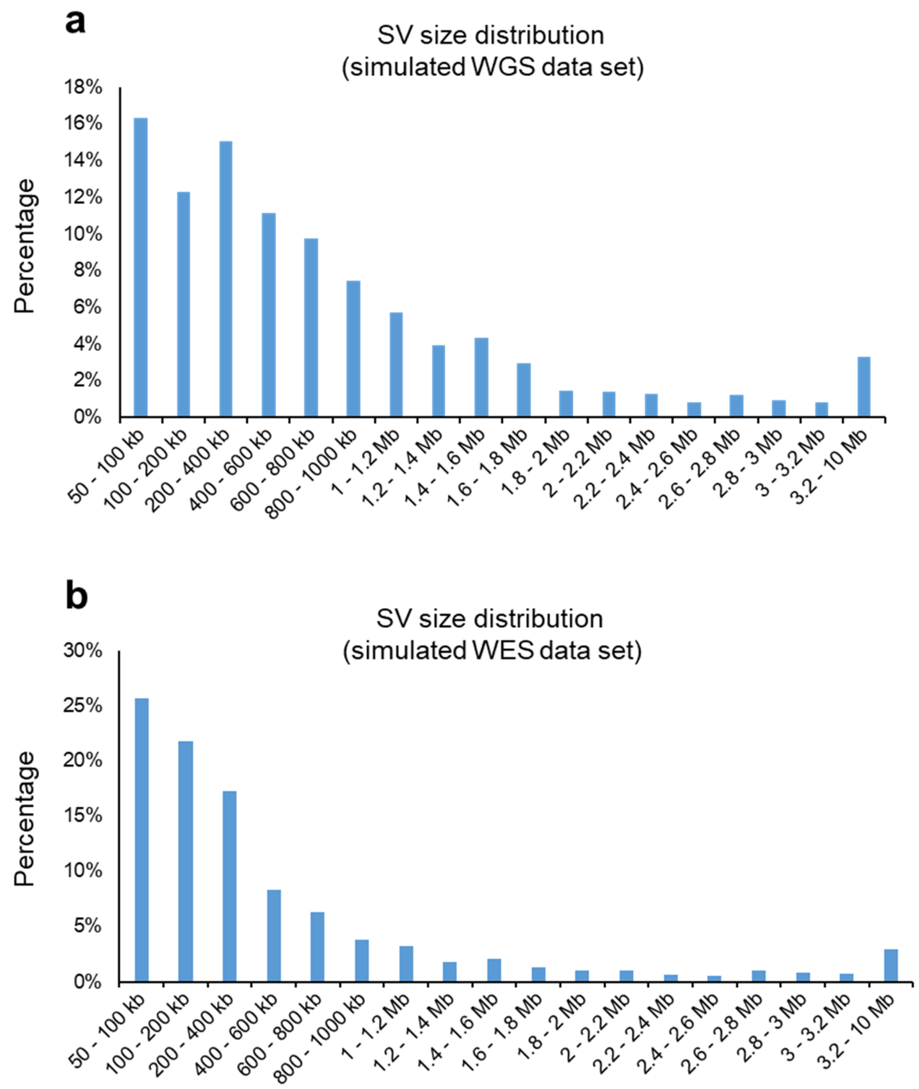


**Supplementary Figure 16**

Size distribution of simulated SV events. **a**) Size distribution of the SVs in the simulated WGS data set. **b**) Size distribution of the SVs in the simulated WES data set. Source data is provided as a Source Data file.

### Supplementary Tables

#### Supplementary Table 1

SV calls detected by Delly on the *F8* inversion sample (10 kb upstream/downstream of the inversion).

| Chrom | Position 1 | Position 2 | SV Type |
|---|---|---|---|
| chrX | 154002063 | 154002446 | INV |
| chrX | 154004508 | 154004709 | INV |
| chrX | 154005670 | 154005855 | INV |
| chrX | 154011709 | 154012062 | DUP |
| chrX | 154012155 | 154012297 | INV |
| chrX | 154014432 | 154014542 | INV |
| chrX | 154014535 | 154014759 | INV |
| chrX | 154017795 | 154017948 | INV |
| chrX | 154020109 | 154020407 | INV |
| chrX | 154045287 | 154045525 | INV |
| chrX | 154045633 | 154045781 | INV |
| chrX | 154057649 | 154057833 | INV |
| chrX | 154057785 | 154057923 | INV |
| chrX | 154083952 | 154084185 | INV |
| chrX | 154091092 | 154091440 | INV |
| chrX | 154115196 | 154115584 | INV |
| chrX | 154115667 | 154115957 | INV |
| chrX | 154124101 | 154124495 | DUP |
| chrX | 154129769 | 154129919 | INV |
| chrX | 154130086 | 154130222 | INV |
| chrX | 154132781 | 154133165 | DUP |
| chrX | 154146261 | 154146462 | INV |
| chrX | 154146565 | 154146775 | INV |
| chrX | 154152621 | 154152772 | INV |
| chrX | 154157646 | 154157941 | DUP |
| chrX | 154158133 | 154158450 | DUP |
| chrX | 154158358 | 154158630 | INV |
| chrX | 154159826 | 154160143 | INV |
| chrX | 154175838 | 154176055 | INV |
| chrX | 154176073 | 154176451 | INV |

| | | | |
|---|---|---|---|
| chrX | 154194502 | 154194721 | INV |
| chrX | 154208905 | 154209847 | DUP |
| chrX | 154209557 | 154209679 | INV |
| chrX | 154212475 | 154212843 | INV |
| chrX | 154212596 | 154212733 | INV |
| chrX | 154227431 | 154227667 | INV |
| chrX | 154272329 | 154272476 | INV |
| chrX | 154272335 | 154272592 | DUP |
| chrX | 154275546 | 154275664 | INV |
| chrX | 154275671 | 154275934 | INV |
| chrX | 154290114 | 154290323 | INV |
| chrX | 154344170 | 154344430 | INV |
| chrX | 154387813 | 154429972 | INV |
| chrX | 154387928 | 154429779 | INV |
| chrX | 154391126 | 154391291 | INV |
| chrX | 154426121 | 154426264 | INV |
| chrX | 154428351 | 154428890 | DUP |
| chrX | 154464614 | 154464716 | INV |
| chrX | 154508570 | 154508865 | INV |
| chrX | 154540689 | 154540872 | INV |
| chrX | 154609922 | 154610441 | DUP |
| chrX | 154610132 | 154610342 | INV |
| chrX | 154611498 | 154611783 | INV |
| chrX | 154611510 | 154688681 | DEL |
| chrX | 154612459 | 154612925 | DUP |
| chrX | 154613017 | 154613356 | INV |
| chrX | 154652935 | 154653319 | DUP |
| chrX | 154687101 | 154687535 | DUP |
| chrX | 154687312 | 154687480 | INV |
| chrX | 154688521 | 154688851 | INV |
| chrX | 154688714 | 154688969 | INV |
| chrX | 154689667 | 154689984 | INV |
| chrX | 154735308 | 154735475 | INV |
| chrX | 154755098 | 154755225 | INV |

**Supplementary Table 2**

SV calls detected by Lumpy on the *F8* inversion sample (all SV calls in chrX).

| Chrom | Position 1 | Position 2 | SV Type |
|---|---|---|---|
| chrX | 1413852 | 1414637 | DEL |
| chrX | 17520630 | 17520931 | DEL |
| chrX | 28749622 | 28749739 | DUP |
| chrX | 31189715 | 31189837 | DUP |
| chrX | 38145097 | 38145387 | DUP |
| chrX | 38145261 | 38145866 | DUP |
| chrX | 38145731 | 38145851 | DEL |
| chrX | 38145513 | 38145876 | DEL |
| chrX | 38145369 | 38145943 | DUP |
| chrX | 38145283 | 38145943 | DUP |
| chrX | 38145180 | 38145975 | DUP |
| chrX | 38145091 | 38145976 | DUP |
| chrX | 38145835 | 38145920 | DEL |
| chrX | 38145729 | 38145981 | DUP |
| chrX | 38145874 | 38145914 | DEL |
| chrX | 38145321 | 38145988 | DUP |
| chrX | 38145471 | 38145995 | DUP |
| chrX | 38145929 | 38146016 | DUP |
| chrX | 38145971 | 38146003 | DEL |
| chrX | 38145885 | 38146041 | DUP |
| chrX | 38145642 | 38146055 | DUP |
| chrX | 38146077 | 38146257 | DUP |
| chrX | 39932139 | 39932332 | DUP |
| chrX | 41788764 | 41788966 | DUP |
| chrX | 47308661 | 47308783 | DUP |
| chrX | 63411862 | 63412017 | DUP |
| chrX | 64063531 | 64063690 | DUP |
| chrX | 76890054 | 76890207 | DUP |
| chrX | 101912168 | 101912314 | DUP |
| chrX | 142794887 | 142795158 | DUP |
| chrX | 149745015 | 149745169 | DUP |

**Supplementary Table 3**

SVs on chrX detected by Longranger on the *F8* inversion sample.

| Chrom 1 | Position 1 | Chrom 2 | Position 2 | SV Type | Distance to int22h-1 | Distance to int22h-3 |
|---|---|---|---|---|---|---|
| chrX | 3735199 | chrX | 3855112 | Unknown | -150380286 | -150832314 |
| chrX | 7810783 | chrX | 8151613 | Unknown | -146304702 | -146535813 |
| chrX | 7810783 | chrX | 8113116 | Unknown | -146304702 | -146574310 |
| chrX | 134855892 | chrX | 134985727 | Unknown | -19259593 | -19701699 |
| chrX | 152225317 | chrX | 152352113 | Unknown | -1890168 | -2335313 |
| chrX | 154091033 | chrX | 154660067 | Unknown | -24452 | -27359 |
| chrX | 154131339 | chrX | 154735755 | Unknown | 15854 | 48329 |

**Supplementary Table 4**

SVs detected by GROC-SVs on the *F8* inversion sample.

| Chrom 1 | Position 1 | Chrom 2 | Position 2 | Orientation |
|---|---|---|---|---|
| chr1 | 12889258 | chr1 | 12938883 | +- |
| chr1 | 12919205 | chr1 | 12853022 | -- |
| chr1 | 21754487 | chr1 | 21794667 | +- |
| chr1 | 148026036 | chr1 | 144622101 | ++ |
| chr1 | 223725671 | chr1 | 223797843 | +- |
| chr2 | 98162357 | chr2 | 97860318 | -- |
| chr2 | 149687145 | chr2 | 149790581 | +- |
| chr2 | 234053740 | chr2 | 234002965 | -+ |
| chr3 | 129809651 | chr3 | 129762840 | -+ |
| chr4 | 9452594 | chr4 | 9485059 | +- |
| chr4 | 69681327 | chr4 | 69893173 | -- |
| chr5 | 155137378 | chr5 | 155188725 | +- |
| chr5 | 180430554 | chr5 | 180375038 | -+ |
| chr6 | 29909733 | chr6 | 29843849 | -- |
| chr6 | 29913575 | chr6 | 29844437 | ++ |
| chr6 | 160956444 | chr6 | 160877743 | -- |
| chr7 | 100550750 | chr7 | 100609409 | -- |
| chr7 | 100555813 | chr7 | 100610572 | ++ |
| chr11 | 1162747 | chr11 | 1212758 | +- |
| chr11 | 5809264 | chr11 | 5777102 | -+ |
| chr12 | 7239875 | chr12 | 7189849 | -+ |
| chr12 | 11545335 | chr12 | 11503243 | -+ |
| chr12 | 18018173 | chr12 | 17922871 | -- |
| chr12 | 109372790 | chr12 | 109423610 | +- |
| chr12 | 132926655 | chr16 | 86452753 | +- |
| chr12 | 133041064 | chr2 | 231869678 | +- |
| chr13 | 114325993 | chr13 | 114425990 | +- |
| chr14 | 24436900 | chr14 | 24474868 | +- |
| chr15 | 20740860 | chr15 | 23406226 | +- |
| chr15 | 22743596 | chr15 | 23572115 | +- |
| chr15 | 23407964 | chr15 | 20739466 | +- |
| chr15 | 23573198 | chr15 | 22742499 | +- |
| chr15 | 28597075 | chr15 | 28806596 | ++ |
| chr15 | 28804860 | chr15 | 28595402 | -- |
| chr15 | 83003956 | chr15 | 82934463 | -- |

| | | | | |
|---|---|---|---|---|
| chr15 | 83014625 | chr15 | 82936703 | ++ |
| chr15 | 84960511 | chr15 | 84859997 | ++ |
| chr16 | 14988609 | chr16 | 15031370 | -- |
| chr16 | 70009791 | chr16 | 74426101 | -+ |
| chr16 | 86453332 | chr12 | 132926283 | +- |
| chr17 | 36297237 | chr17 | 36337601 | +- |
| chr20 | 1600172 | chr20 | 1559471 | -+ |
| chr20 | 56771729 | chr12 | 2828954 | -- |
| chr20 | 56771963 | chr12 | 2830313 | ++ |
| chr22 | 18666166 | chr22 | 18737108 | -- |
| chr22 | 18737933 | chr22 | 18686213 | ++ |

**Supplementary Table 5**

SVs on chrX detected by NAIBR on the *F8* inversion sample.

| Chrom 1 | Position 1 | Chrom 2 | Position 2 | Orientation | Distance to int22h-1 | Distance to int22h-3 |
|---|---|---|---|---|---|---|
| chrX | 1399792 | chrX | 1400312 | +- | -152715693 | -153287114 |
| chrX | 26179787 | chrX | 26212951 | ++ | -127935698 | -128474475 |
| chrX | 49162006 | chrX | 49180528 | ++ | -104953479 | -105506898 |
| chrX | 49208629 | chrX | 49209218 | +- | -104906856 | -105478208 |
| chrX | 49218203 | chrX | 49218751 | +- | -104897282 | -105468675 |
| chrX | 52830446 | chrX | 52830826 | +- | -101285039 | -101856600 |
| chrX | 57147470 | chrX | 57162963 | ++ | -96968015 | -97524463 |
| chrX | 129651865 | chrX | 129652250 | +- | -24463620 | -25035176 |
| chrX | 140140043 | chrX | 140140427 | +- | -13975442 | -14546999 |
| chrX | 154387911 | chrX | 154430070 | -- | 272426 | -257356 |
| chrX | 155245001 | chrX | 155245507 | +- | 1129516 | 558081 |
| chrX | 155251108 | chrX | 155252593 | +- | 1135623 | 565167 |

**Supplementary Table 6**

SVs on chr17 detected by NAIBR on the *NF1* sample.

| Chrom 1 | Position 1 | Chrom 2 | Position 2 | Orientation | Distance to breakpoint 1 | Distance to breakpoint 2 |
|---|---|---|---|---|---|---|
| chr17 | 10359246 | chr17 | 10409257 | -- | -19325000 | -19413270 |
| chr17 | 18296487 | chr17 | 18296856 | +- | -11387759 | -11525671 |
| chr17 | 18314761 | chr17 | 18315123 | +- | -11369485 | -11507404 |
| chr17 | 18327938 | chr17 | 18328330 | +- | -11356308 | -11494197 |
| chr17 | 18343774 | chr17 | 18344042 | +- | -11340472 | -11478485 |
| chr17 | 18390522 | chr17 | 18391011 | +- | -11293724 | -11431516 |
| chr17 | 27963239 | chr17 | 27964080 | +- | -1721007 | -1858447 |
| chr17 | 34502325 | chr17 | 34502718 | +- | 4818079 | 4680191 |
| chr17 | 39241002 | chr17 | 39296331 | -+ | 9556756 | 9473804 |
| chr17 | 39241014 | chr17 | 39274415 | -- | 9556768 | 9451888 |
| chr17 | 39254370 | chr17 | 39279917 | -+ | 9570124 | 9457390 |
| chr17 | 39382979 | chr17 | 39411518 | -+ | 9698733 | 9588991 |
| chr17 | 39421637 | chr17 | 39427775 | +- | 9737391 | 9605248 |
| chr17 | 39421677 | chr17 | 39432260 | +- | 9737431 | 9609733 |
| chr17 | 39502590 | chr17 | 39521159 | -- | 9818344 | 9698632 |
| chr17 | 40653142 | chr17 | 40697232 | ++ | 10968896 | 10874705 |
| chr17 | 41008335 | chr17 | 41026393 | ++ | 11324089 | 11203866 |
| chr17 | 43616656 | chr17 | 43616962 | +- | 13932410 | 13794435 |
| chr17 | 58073089 | chr17 | 58073319 | +- | 28388843 | 28250792 |
| chr17 | 58078460 | chr17 | 58078688 | +- | 28394214 | 28256161 |
| chr17 | 60343141 | chr17 | 60343406 | +- | 30658895 | 30520879 |
| chr17 | 60359848 | chr17 | 60360239 | +- | 30675602 | 30537712 |
| chr17 | 62213073 | chr17 | 62214273 | +- | 32528827 | 32391746 |
| chr17 | 62885755 | chr17 | 62886258 | +- | 33201509 | 33063731 |
| chr17 | 64794832 | chr17 | 64795449 | +- | 35110586 | 34972922 |
| chr17 | 78287778 | chr17 | 78289508 | +- | 48603532 | 48466981 |
| chr17 | 80105167 | chr17 | 80106015 | +- | 50420921 | 50283488 |

**Supplementary Table 7**

SV calls detected by Delly on the *NF1* deletion sample (1 Mb upstream/downstream of the deletion).

| Chrom | Position 1 | Position 1 | SV Type | SV Length |
|---|---|---|---|---|
| chr17 | 17363396 | 79498375 | INV | 62134980 |
| chr17 | 17363532 | 64795530 | INV | 47431999 |
| chr17 | 17363628 | 79499683 | INV | 62136056 |
| chr17 | 18286739 | 74017685 | INV | 55730947 |
| chr17 | 18286866 | 74017502 | INV | 55730637 |
| chr17 | 28749720 | 28749884 | INV | 165 |
| chr17 | 28749894 | 28750067 | INV | 174 |
| chr17 | 28778475 | 28778674 | INV | 200 |
| chr17 | 28778804 | 28778982 | INV | 179 |
| chr17 | 28781391 | 28781646 | INV | 256 |
| chr17 | 28789399 | 28789573 | INV | 175 |
| chr17 | 28804180 | 28804452 | INV | 273 |
| chr17 | 28850972 | 28851083 | INV | 112 |
| chr17 | 28880548 | 28880795 | INV | 248 |
| chr17 | 28883413 | 28883543 | INV | 131 |
| chr17 | 28883874 | 28884185 | DUP | 312 |
| chr17 | 28885891 | 28885977 | INV | 87 |
| chr17 | 28886217 | 28886887 | DUP | 671 |
| chr17 | 28886790 | 28887002 | INV | 213 |
| chr17 | 28896496 | 28896657 | INV | 162 |
| chr17 | 28898325 | 28898507 | INV | 183 |
| chr17 | 28898453 | 28898776 | INV | 324 |
| chr17 | 28900688 | 28900885 | INV | 198 |
| chr17 | 28958756 | 28958860 | INV | 105 |
| chr17 | 28963620 | 28963782 | INV | 163 |
| chr17 | 29161159 | 29161443 | INV | 285 |
| chr17 | 29214324 | 29214454 | INV | 131 |
| chr17 | 29233308 | 29233456 | INV | 149 |
| chr17 | 29235695 | 29235926 | INV | 232 |
| chr17 | 29272077 | 29272279 | INV | 203 |
| chr17 | 29280208 | 29280325 | INV | 118 |
| chr17 | 29280249 | 29280542 | DUP | 294 |
| chr17 | 29292652 | 29292783 | INV | 132 |

| | | | | |
|---|---|---|---|---|
| chr17 | 29297874 | 29298071 | INV | 198 |
| chr17 | 29311663 | 29311773 | INV | 111 |
| chr17 | 29312096 | 29312227 | INV | 132 |
| chr17 | 29323985 | 29324271 | INV | 287 |
| chr17 | 29328060 | 29328325 | INV | 266 |
| chr17 | 29374344 | 29374472 | INV | 129 |
| chr17 | 29375112 | 29375224 | INV | 113 |
| chr17 | 29375965 | 29376167 | INV | 203 |
| chr17 | 29376169 | 29376747 | DUP | 579 |
| chr17 | 29376450 | 29376678 | INV | 229 |
| chr17 | 29376869 | 29377109 | INV | 241 |
| chr17 | 29377045 | 29377508 | DUP | 464 |
| chr17 | 29377115 | 29377278 | INV | 164 |
| chr17 | 29422253 | 29422388 | INV | 136 |
| chr17 | 29442254 | 29442686 | DUP | 433 |
| chr17 | 29442303 | 29442458 | INV | 156 |
| chr17 | 29482195 | 29482490 | INV | 296 |
| chr17 | 29496377 | 29496593 | INV | 217 |
| chr17 | 29499399 | 29499499 | INV | 101 |
| chr17 | 29499572 | 29499701 | INV | 130 |
| chr17 | 29509791 | 29509902 | INV | 112 |
| chr17 | 29553393 | 29553609 | INV | 217 |
| chr17 | 29559956 | 29560152 | INV | 197 |
| chr17 | 29631993 | 29632186 | INV | 194 |
| chr17 | 29646082 | 29646292 | INV | 211 |
| chr17 | 29653050 | 29653191 | INV | 142 |
| chr17 | 29653320 | 29653427 | INV | 108 |
| chr17 | 29670886 | 29671266 | INV | 381 |
| chr17 | 29688359 | 29688466 | INV | 108 |
| chr17 | 29691368 | 29691611 | INV | 244 |
| chr17 | 29792500 | 29792713 | INV | 214 |
| chr17 | 29844524 | 29845097 | DUP | 574 |
| chr17 | 29844528 | 29844681 | INV | 154 |
| chr17 | 29844742 | 29844865 | INV | 124 |
| chr17 | 29848528 | 29848642 | INV | 115 |
| chr17 | 29848670 | 29848862 | INV | 193 |
| chr17 | 29897228 | 29897443 | INV | 216 |
| chr17 | 29897946 | 29898219 | INV | 274 |
| chr17 | 29898357 | 29898503 | INV | 147 |

| | | | | |
|---|---|---|---|---|
| chr17 | 30032123 | 30032285 | INV | 163 |
| chr17 | 30032425 | 30032534 | INV | 110 |
| chr17 | 30098240 | 30098400 | INV | 161 |
| chr17 | 30185676 | 30185833 | INV | 158 |
| chr17 | 30262037 | 30262472 | DUP | 436 |
| chr17 | 30315050 | 30315340 | INV | 291 |
| chr17 | 30321519 | 30321682 | INV | 164 |
| chr17 | 30347576 | 30348121 | INV | 546 |
| chr17 | 30348192 | 30348503 | INV | 312 |
| chr17 | 30348913 | 30349157 | INV | 245 |
| chr17 | 30362642 | 30362791 | INV | 150 |
| chr17 | 30411530 | 30411734 | INV | 205 |
| chr17 | 30418743 | 30418959 | INV | 217 |
| chr17 | 30477548 | 30477908 | INV | 361 |
| chr17 | 30537828 | 30538161 | DUP | 334 |
| chr17 | 30537980 | 30538161 | INV | 182 |
| chr17 | 30594987 | 30595297 | INV | 311 |
| chr17 | 30601399 | 30601626 | INV | 228 |
| chr17 | 30615745 | 30615901 | INV | 157 |
| chr17 | 30662014 | 30662173 | INV | 160 |
| chr17 | 30689981 | 30690196 | INV | 216 |
| chr17 | 30692181 | 30692341 | INV | 161 |
| chr17 | 30692457 | 30692615 | INV | 159 |
| chr17 | 30693806 | 30693985 | INV | 180 |
| chr17 | 30694745 | 30694950 | INV | 206 |
| chr17 | 30771326 | 30771549 | INV | 224 |
| chr17 | 30800970 | 30801130 | INV | 161 |
| chr17 | 30806902 | 30807104 | INV | 203 |
| chr17 | 30806952 | 30807125 | INV | 174 |
| chr17 | 30822023 | 30822151 | INV | 129 |

**Supplementary Table 8**

SV calls detected by Lumpy on the *NF1* deletion sample (1 Mbp upstream/downstream of the deletion).

| Chrom | Position 1 | Position 1 | SV Type | SV Length |
|---|---|---|---|---|
| chr17 | 28749892 | 28750050 | INV | 158 |
| chr17 | 28804393 | 28804502 | INV | 109 |
| chr17 | 28811314 | 28811822 | INV | 508 |
| chr17 | 28883873 | 28884184 | DUP | 311 |
| chr17 | 28886443 | 28886674 | INV | 231 |
| chr17 | 28886217 | 28886894 | DUP | 677 |
| chr17 | 28886812 | 28887273 | INV | 461 |
| chr17 | 28898494 | 28898775 | INV | 281 |
| chr17 | 29280248 | 29280541 | DUP | 293 |
| chr17 | 29312021 | 29312222 | INV | 201 |
| chr17 | 29376217 | 29376575 | INV | 358 |
| chr17 | 29376197 | 29376860 | DUP | 663 |
| chr17 | 29377124 | 29377287 | INV | 163 |
| chr17 | 29377044 | 29377507 | DUP | 463 |
| chr17 | 29422255 | 29422415 | INV | 160 |
| chr17 | 29442358 | 29442535 | INV | 177 |
| chr17 | 29499570 | 29499688 | INV | 118 |
| chr17 | 29553392 | 29553609 | INV | 217 |
| chr17 | 29559812 | 29559979 | INV | 167 |
| chr17 | 29585386 | 29585478 | INV | 92 |
| chr17 | 29631907 | 29632107 | INV | 200 |
| chr17 | 29677233 | 29677432 | INV | 199 |
| chr17 | 29691366 | 29691580 | INV | 214 |
| chr17 | 29844728 | 29844864 | INV | 136 |
| chr17 | 29844523 | 29844898 | DUP | 375 |
| chr17 | 29848568 | 29848854 | INV | 286 |
| chr17 | 30594805 | 30594919 | INV | 114 |
| chr17 | 30594966 | 30595296 | BND | 330 |
| chr17 | 30692180 | 30692359 | INV | 179 |
| chr17 | 30806472 | 30806667 | INV | 195 |
| chr17 | 30806578 | 30807234 | DUP | 656 |
| chr17 | 18286866 | 74017503 | BND | 55730637 |
| chr17 | 7167959 | 30145402 | BND | 22977443 |

**Supplementary Table 9**

SVs detected by GROC-SVs on the *NF1* WES sample (on chr17)

| Chrom 1 | Position 1 | Chrom 2 | Position 2 | Orientation |
|---|---|---|---|---|
| chr17 | 397512 | chr17 | 287512 | -+ |
| chr17 | 287512 | chr17 | 397512 | +- |

**Supplementary Table 10**

Simulated CNVs that are known to cause CNV syndromes (related to Supplementary Figure 7)

| Chrom | Position 1 | Position 1 | SV Type | SV Length |
|---|---|---|---|---|
| chr1 | 10001 | 12840260 | DEL | 12830259 |
| chr1 | 145386506 | 145748068 | DEL | 361562 |
| chr1 | 146533376 | 147883377 | DUP | 1350001 |
| chr2 | 44410451 | 44589585 | DEL | 179134 |
| chr2 | 59285696 | 61819816 | DEL | 2534120 |
| chr2 | 196925121 | 205206940 | DEL | 8281819 |
| chr2 | 239969863 | 240322644 | DEL | 352781 |
| chr3 | 195726835 | 197344664 | DUP | 1617829 |
| chr4 | 1569197 | 2110237 | DEL | 541040 |
| chr5 | 10001 | 12533305 | DEL | 12523304 |
| chr5 | 112043201 | 112181937 | DEL | 138736 |
| chr5 | 126112314 | 126172713 | DUP | 60399 |
| chr5 | 175724636 | 177052117 | DEL | 1327481 |
| chr7 | 72744455 | 74142673 | DUP | 1398218 |
| chr7 | 96318078 | 96339204 | DEL | 21126 |
| chr8 | 8100055 | 11764630 | DUP | 3664575 |
| chr8 | 77226464 | 77766240 | DEL | 539776 |
| chr9 | 140513443 | 140730579 | DEL | 217136 |
| chr11 | 31806339 | 32457088 | DEL | 650749 |
| chr11 | 43994800 | 46052451 | DEL | 2057651 |
| chr12 | 1080000 | 1346472 | DEL | 266472 |
| chr12 | 65071919 | 68645526 | DEL | 3573607 |
| chr15 | 22749354 | 28438267 | DEL | 5688913 |
| chr15 | 30910306 | 32445408 | DEL | 1535102 |
| chr15 | 74412643 | 75972912 | DEL | 1560269 |
| chr15 | 99357970 | 102521393 | DUP | 3163423 |
| chr16 | 60001 | 834373 | DEL | 774372 |

| | | | | |
|---|---|---|---|---|
| chr16 | 3775055 | 3930122 | DEL | 155067 |
| chr16 | 14986684 | 16486685 | DUP | 1500001 |
| chr16 | 21475060 | 29284078 | DUP | 7809018 |
| chr16 | 29606852 | 30199856 | DUP | 593004 |
| chr17 | 1 | 2588910 | DEL | 2588909 |
| chr17 | 14097915 | 15470904 | DUP | 1372989 |
| chr17 | 16773072 | 20222150 | DUP | 3449078 |
| chr17 | 29107097 | 30263322 | DEL | 1156225 |
| chr17 | 34815072 | 36215918 | DEL | 1400846 |
| chr17 | 43705166 | 44294407 | DEL | 589241 |
| chr21 | 27252860 | 27543447 | DUP | 290587 |
| chr22 | 1 | 16971861 | DUP | 16971860 |
| chr22 | 19009792 | 21452446 | DEL | 2442654 |
| chr22 | 21917117 | 23722446 | DEL | 1805329 |
| chr22 | 51045516 | 51187845 | DEL | 142329 |
| chrX | 751878 | 867876 | DEL | 115998 |
| chrX | 6455812 | 8133196 | DEL | 1677384 |
| chrX | 48334549 | 52117662 | DUP | 3783113 |
| chrX | 53401070 | 53683276 | DUP | 282206 |
| chrX | 103031438 | 103047548 | DUP | 16110 |
| chrX | 153287263 | 153363189 | DUP | 75926 |
| chrX | 153624563 | 153881854 | DUP | 257291 |
| chrY | 14352761 | 15154863 | DEL | 802102 |
| chrY | 20118045 | 26065198 | DEL | 5947153 |

#### Supplementary Methods

##### Detection of SVs using different SV callers

SV detection using Longranger (version 2.2.2) was performed with default settings for detection of germline SVs, and the "--somatic" parameter was set for detection of SVs with VAF of 10% and 20%. SV detection using GROC-SVs (version 0.2.5) was performed using default settings with a blacklist file from 10X Genomics. SV detection using NAIBR (commit 15eba96) was performed using default settings a blacklist file from 10X Genomics. SV detection using Delly (version 0.8.1) was performed with default settings. SV detection using Lumpy (version 0.3.0) was performed using the smoove pipeline (https://github.com/brentp/smoove) according to the authors' suggestions. *De novo* assembly based SV calling using FermiKit (commit bf9c711) was performed with the parameters "-s 3g -l 151" to specify the expected genome size and read length. Detection of SVs from the HX1 PacBio data set using Sniffles (version 1.0.11) was performed with the "--min_support 1" parameter to set the minimum number of reads that support an SV to be 1. The purpose is to maximize the sensitivity and see if Sniffles can detect the duplications reported by SMRT-SV.

##### Error correction of PacBio reads

PacBio sequencing data of the HX1 genome was downloaded from NCBI SRA database with the accession SRX1424851. Raw PacBio reads (subreads) were extracted from the HDF5 files using pbh5tools according to the manufacturer's instructions. Error correction of PacBio reads was performed using Canu[1] (version 1.6) with the parameters for PacBio reads (canu -correct genomeSize=3.1G -pacbio-raw path/to/pacbio_reads.fasta).

**Mapping of PacBio data to the human genome**

Raw PacBio reads or error-corrected PacBio reads were mapped to human reference genome GRCh38 (without ALT loci) using Minimap2[2] (version: 2.15-r905) with parameters optimized for PacBio reads (-ax map-pb). The output SAM file was converted to BAM file and sorted using SAMtools[3].

**De novo assembly of the PacBio reads supporting the SV**

SAMtools was used to extract the PacBio reads that were aligned to the proposed variant allele in HX1 (related to Supplementary Figure 9). *De novo* assembly of the extracted reads was performed using wtdbg2[4] with the parameters "-x rsII -g 50k" to specify that the platform was PacBio RS II and the estimated size of the target region was 50 kb. A single contig of 42.7 kb was generated.

**Comparison of the SV call sets of LinkedSV and SMRT-SV on the HX1 genome**

The SV calls of SMRT-SV were downloaded from NCBI dbVar database under accession nstd162. We only compared large SV calls that are at least 10 kb, because LinkedSV had limited power to detect smaller SVs except small deletions. The performance of small deletion detection was evaluated on the GIAB HG002 genome. Two SV calls were considered the same if they had at least 50% reciprocal overlap (the overlapped region was more than 50% of both calls). This criterion was chosen to follow what was done by a previous study [5].

#### Supplementary Notes

**Supplementary Note 1: Explanation of the model for detecting type 2 evidence.**

Barcode similarity between two nearby regions very high because the reads originate from almost the same set of HMW DNA molecules. However, the barcode similarity between the left side and right side of the breakpoint are dramatically reduced. We call this evidence as type 2 evidence. To detect type 2 evidence, LinkedSV uses two adjacent sliding windows (window 1 and window 2) to scan the genome and calculate the barcode similarity between the window 1 and window 2.

In WES data sets, the numbers of reads in the sliding windows vary a lot due to capture bias and the length of capture regions. To detect type 2 evidence from both WGS and WES data sets, our model considers the variation of sequencing depth and capture regions. The barcode similarity is calculated as:

$$S = \frac{x}{m_1^a m_2^b} n e^{-\alpha d} \quad (1),$$

where:

$m_1$ is the number of barcodes in window 1,

$m_2$ is the number of barcodes in window 2,

$x$ is the number of barcodes in both windows,

$d$ is the weight distance between reads of the left window and the right window,

$n$ is a constant representing the characteristic of the library,

$\alpha$ is a parameter of fragment length distribution,

a and b are two parameters between 0 and 1,

$n$, $\alpha$, $a$ and $b$ are estimated from the data using regression.

Suppose there are $n$ different HMW DNA molecules span both window 1 and window 2, each of which has a different barcode and generates a number of read pairs in the library. The read pairs in the library may or may not be sequenced. We assume the $n$ HMW DNA molecules have the same rate to generate read pairs in the library, so the $n$ HMW DNA molecules have the same chance to be sequenced (have at least 1 read). Let $m_1$ be the number of HMW DNA molecules sequenced in window 1, $m_2$ be the number of HMW DNA molecules sequenced in window 2. $m_1$ and $m_2$ can be different due to the bias of target enrichment and the total length of target regions in each window. Let $X$ be the number of HMW DNA molecules sequenced in both window 1 and window 2. $X$ follows the hypergeometric distribution:

$$P(X = x \ |m_1, m_2, n) = \frac{C_{m_1}^{x} C_{n-m_1}^{m_2-x}}{C_n^{m_2}} \quad (2).$$

The expectation of X is:

$$E(X) = \frac{m_1 m_2}{n} \ (3).$$

However, the length of sliding windows may be as long as 40 kb and not all the $n$ HMW DNA molecules are long enough to span both windows. In addition, the capture regions in window 1 and window 2 may be close to each other or far away from each other. Therefore, we need to adjust $n$ to be approximately $ne^{-\alpha d}$. $d$ is calculated using the following equation:

$d = w_2 - w_1$ (4),

where $w_1$ is the mean mapping position of all reads in window 1 and $w_2$ is the mean mapping position of all reads in window 2. The larger $d$, the smaller number of HMW DNA molecules can span a region of length $d$. We choose exponential distribution because the length of HMW DNA molecules follows exponential distribution and thus the number of HMW DNA molecules longer than $d$ also follows exponential distribution.

$m_1$ also need to be adjusted because not all HMW DNA molecules being sequenced in window 1 span both windows. We adjust $m_1$ to be approximately $m_1^a$ and similarly adjust $m_2$ to be approximately $m_2^b$.

After the adjustment, the expectation of $X$ is:

$E(X) = \frac{m_1^a m_2^b}{ne^{-\alpha d}}$ (5).

We define barcode similarity as:

$S = \frac{x}{E(X)} = \frac{x}{m_1^a m_2^b} ne^{-\alpha d}$ (6),

where x is the number of shared barcodes between window 1 and 2, $E(X)$ is the expected number of shared barcodes between window 1 and 2.

Take the log of both sides equation (6), we have:

$\log(S) = \log(x) - alog(m_1) - blog(m_2) + \log(n) - \alpha d$ (7).

Assuming most regions in the genome do not have breakpoints, we can replace $S$ with 1 and estimate $a$, $b$, $n$, $\alpha$ from the data using linear regression.

**Supplementary Note 2: Range of SV size that can be detected by LinkedSV**

LinkedSV is able to detect deletions ≥ 50 bp, inversions ≥ 10 kb, tandem duplications ≥ 20 kb and intra-chromosomal translocations of any size. Supplementary Figure 8 showed the size distribution of SV events detected from the HG002 genome using LinkedSV.

LinkedSV has limited power to detect small SV duplications and inversions. Fortunately, a lot of large SVs are known to be associated with human diseases. We analyzed the sizes of SVs from the following two resources:

1) Somatic Cancer SVs released by the COSMIC database [6] (version 89, released May 15$^{th}$, 2019). This data set contains 351,862 SVs.

2) Expert-curated known SVs that cause CNV syndromes. The DECIPHER (Database of Chromosomal Imbalance and Phenotype in Humans Using Ensembl Resources) database[7] provides a list of expert-curated microdeletion and microduplication syndromes involved in developmental disorders. The genomic coordinates of the CNVs were obtained from https://decipher.sanger.ac.uk/disorders#syndromes/overview . This data set contains 67 SVs.

The SV size distribution of the two data sets were shown in Supplementary Figure 5. 71% of the cancer somatic SVs released by COSMIC database are inter-chromosomal events or intra-chromosomal events that are larger than 10 kb. All deletions/duplications that cause the CNV syndromes are larger than 10 kb. The results indicated that large SVs (including inter-chromosomal SVs) are associated with diseases such as cancers and CNV syndromes. Of note, we need to be aware that there is potential bias in calculating the fraction of large CNVs, as large CNVs are easier to detect and there may exist disease-associated small CNVs that have not been detected.
